# Cell-free synthesis of natural compounds from genomic DNA of biosynthetic gene clusters

**DOI:** 10.1101/2020.04.04.025353

**Authors:** Ilka Siebels, Sarah Nowak, Christina S. Heil, Peter Tufar, Niña S. Cortina, Helge B. Bode, Martin Grininger

## Abstract

A variety of chemicals can be produced in a living host cell via optimized and engineered biosynthetic pathways. Despite the successes, pathway engineering remains demanding and partly impossible owing to the lack of specific functions or substrates in the host cell, its sensitivity in vital physiological processes to the heterologous components, or constrained mass transfer across the membrane. In this study, we demonstrate that cell-free systems can be useful in driving the characterization and engineering of biosynthetic pathways. We show that complex multidomain proteins involved in natural compound biosynthesis can be produced from encoding DNA *in vitro* in a minimal complex PURE system to directly run multistep reactions. We prove the concept of this approach on the direct synthesis of indigoidine and rhabdopeptides with the *in vitro* produced multidomain megasynthases BpsA and KJ12ABC. The *in vitro* produced proteins are analyzed in detail, i.e., in yield, quality, post-translational modification and specific activity, and compared to recombinantly produced proteins. Our study highlights cell-free PURE systems as suitable setting for the rapid engineering of biosynthetic pathways.

## Introduction

Genome mining has become the main driver in the discovery of new natural products ^1-2^. However, while the “orphan clusters” become available by the advent of next generation sequencing technologies and bioinformatics tools, the substances can often neither be identified nor extracted under laboratory conditions. Cultivation of the native producer strain in the laboratory is of limited success, because the native conditions cannot be reconstituted in the lab ^3^, and robust heterologous production of the compound in host organisms fails owing to complex native regulation and possible toxic products ^4-5^. Concomitantly, the proteins encoded by gene clusters escape enzymatic characterization, catalytic mechanisms remain unsolved and protein engineering for expanding the product spectra to novel new-to-nature compounds remains unachievable.

Facing this problem, we sought to establish a cell-free/*ex vivo* approach based on the PURE system for analyzing and engineering biosynthetic gene clusters. Such an approach is beneficial for the analysis and design of biosynthetic pathways for mainly three reasons: First, it prevents the biosynthetic pathway from interfering with physiological processes of a host cell, second, it is of minimal complexity allowing for simple and direct read-out of enzymatic properties, and, third, it is an open system that is not constrained to canonical compounds (substrates, cofactors, or assisting and tailoring enzymes). Overall, a minimal complex cell-free system could improve access to the key information about native and engineered pathways. In evaluating the PURE system for the cell-free synthesis of natural compounds from genomic DNA, we worked with the commercially available *E. coli*-based PURExpress *In Vitro* Protein Synthesis (IVPS) Kit as a “reaction solution” for gene expression and product formation (New England Biolabs, USA) ^6^. We posited that the *E. coli*-based system provides a suited setting, because a multitude of proteins from biosynthetic gene clusters were successfully produced in *E. coli* ^7-8^.

In our approach, we specifically focused on the protein class of megasynthases, which are responsible for the synthesis of non-ribosomal peptides (NRPs) and polyketides (PKs) ^9-11^. The megasynthases, termed NRP synthetases (NRPSs) and PK synthases (PKSs), are organized in modules comprising integrated enzymatic domains that catalyze the individual reaction steps in biosynthesis ^12-13^. PKSs and NRPSs can occur as monomodular or multimodular systems, while the genes encoding multimodular megasynthases typically cluster in the genome. Intriguingly, there is co-linearity between the order of the genes encoding modules of a multimodular megasynthase, the sequence of the reaction steps assembling the compound, and the identity of the product. Megasynthase gene clusters are prototypical candidates for identification by genome mining, because their inherent modularity leads to gene clusters with repeating patterns. Further, the paradigm of co-linearity leads to accurate predictions about the identity of the compound ^14-15^, which has put megasynthases at the forefront of engineering machineries for the programmable multistep synthesis of novel compounds with new bioactivities ^16-18^. Due to the relevance of PKs and NRPs as pharmaceuticals, a simple and fast access to the products of native and engineered megasynthases is highly demanded.

In this report, we demonstrate the cell-free production of the NRPS BpsA from *Streptomyces lavendulae* ^19-20^ and the RXP (rhabdopeptide-like peptide) producing NRPS KJ12ABC from *Xenorhabdus* KJ12.1 ^21^ in the PURE system, as well as the direct production of their natural products indigoidine and rhabdopeptides, respectively (Figure 1A & B). We further show that other megasynthases (including the PKS-related fatty acid synthases (FASs)) can be produced and activated by post-translational modification. The successful translation of the selected genes into natural products proves the general applicability of PURE systems as cell-free platform for studying megasynthases and biosynthesis of their natural products. The presented approach can be directly applied to other proteins involved in natural compound synthesis.

**Figure 1.**
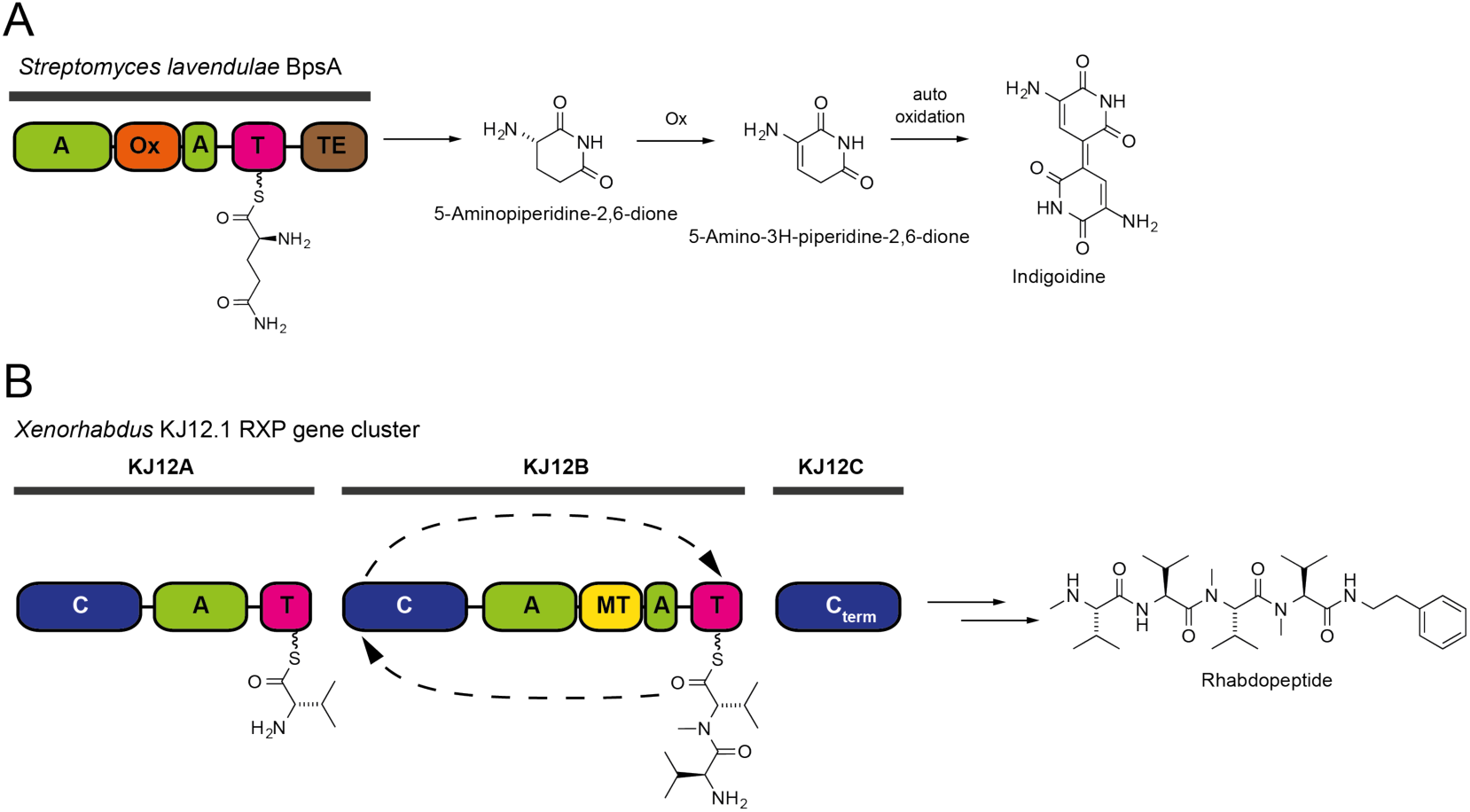
NRPS-mediated indigoidine and rhabdopeptide synthesis. NRPS modules compose three core domains: a condensation (C) domain, an adenylation (A) domain, and a thiolation (T) domain. Additional domains release the compound and may be present for further processing. **(A)** BpsA-mediated indigoidine biosynthesis. The A domain incorporates L-glutamine, which then undergoes internal cyclization. The product is released by a thioesterase (TE) domain. The oxidation (Ox) domain presumably oxidizes the cyclized product. Further oxidation dimerizes the intermediate to form the blue pigment indigoidine ^22^. **(B)** The RXP-producing NRPS from *Xhenorhabdus* KJ12.1, termed KJ12ABC. The RXP biosynthetic gene cluster encodes three NRPS modules. The stand-alone C_term_ domain uses phenylethylamine (PEA) for peptide chain release. Valine is the preferred substrate of both elongation modules KJ12A and KJ12B. The second module KJ12B is iteratively used in this assembly line with a relaxed methyltransferase (MT) activity. KJ12ABC produces a broad spectrum of compounds ^21^.

## Results & Discussion

To establish an *in vitro* platform for megasynthase production and biosynthesis, the monomodular NRPS protein BpsA was produced *in vitro* using the commercially available PURExpress *In Vitro* Protein Synthesis Kit (New England Biolabs, USA) ^6^. This IVPS system is reconstituted from pure components in defined concentrations and based on the *E. coli* transcription/translation machinery. BpsA served as main model system in our study, because it is a monomodular NRPS and produces the blue colored pigment indigoidine that allows the direct monitoring of product synthesis by spectroscopic means. Full-length BpsA (Uniprot number Q1MWN4, N-terminal Strep-Tag ^23^) was successfully synthesized *in vitro*, as confirmed by correct molecular weight bands in SDS-PAGE and by Western blotting (Figure 2A). Size exclusion chromatography (SEC) showed that *in vitro* synthesized BpsA elutes at identical apparent mass as the recombinantly produced reference (Figure 2B). Post-translational phosphopantetheinylation of the thiolation (T) domain, an obligatory modification for enabling substrate shuttling, was performed by the promiscuous 4’-phosphopantetheinyl transferase enzyme Sfp from *Bacillus subtilis* and coenzyme A (CoA) ^24-25^, which were both added to the solution. Protein yields after 2 h of synthesis at 30 °C were in average 25.9 ± 4.0 µg/mL, when the protein was first synthesized and then phosphopantetheinylated (sequential reaction), and 15.3 ± 2.4 µg/mL, when the protein was phosphopantetheinylated while synthesized (parallel reaction) (Figure 2C and Figure S1). We have not further investigated the origin of the difference in expression yields in the sequential and the parallel protocol.

**Figure 2.**
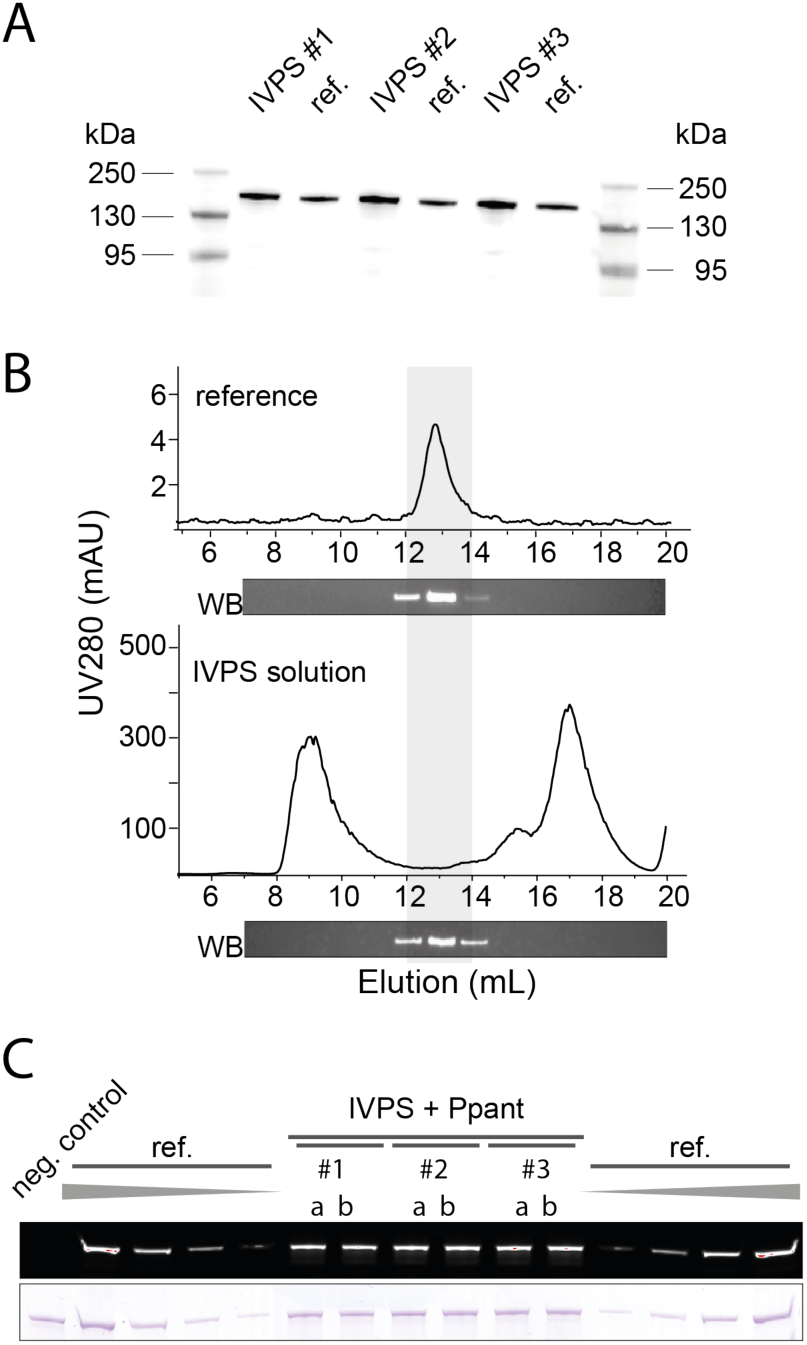
Synthesis of BpsA with the PURE cell-free system. **(A)** Expression control by Western blotting with anti-Strep antibodies performed in three independent reaction solutions (#1-3). BpsA was applied as *holo*-protein, produced by IVPS with simultaneous phosphopantetheinylation. Self-cast 9 % Tris-Tricine gel. Strep-tagged BpsA has a molecular weight of 142.7 kDa. For the uncropped blot, see Figure S2A. **(B)** SEC profiles and Western Blot detection of elution fractions. (top) Recombinantly produced BpsA and (bottom) IVPS reaction solution including phosphopantetheinylation. **(C)** Quantification of protein production yields and phosphopantetheinylation efficiency. BpsA was first produced by IVPS and then phosphopantetheinylated with Sfp and CoA-647 (purchased from NEB). Samples from three independent reactions (#1-3) were applied in repetition (a & b). For calibration, recombinantly produced BpsA, diluted in the PURExpress reaction solution, was loaded in amounts of 1.25, 0.63, 0.31 and 0.16 pmol. 9 % Tris-Tricine gel as in panel A. For the uncropped gels, see Figure S2B. Overall, three times three reactions, each applied in duplicate (18 bands), were used for quantification of BpsA production and phosphopantetheinylation for the parallel and the sequential protocol, respectively (Figure S3 A-C).

The susceptibility of *in vitro* synthesized BpsA for phosphopantetheinylation was determined by labeling with the fluorescent CoA-647 and in-gel fluorescence read-out ^26^. We assumed that the degree of labeling (DOL) with CoA-647 is a good measure for the phosphopantetheinylation efficiency, because Sfp is tolerant for CoA and CoA-derivatives ^27-28^. We found DOLs of 108.1 ± 8.3 % and 127.2 ± 8.4 % for the sequential and the parallel protocol, respectively, when compared to recombinantly produced BpsA as reference (see Figure 2C). DOLs for the *in vitro* synthesized BpsA exceeding 100 % may be explained by the *in vivo* phosphopantetheinylation of a small fraction of the recombinantly produced BpsA that was used as a reference. The phosphopantetheinylated fraction is withdrawn from CoA-647 labeling, leading to the systematic overestimation of DOLs of cell-free produced BpsA.

Subsequently, *in vitro* synthesized and phosphopantetheinylated BpsA was used to biosynthesize indigoidine *in situ*. To initiate indigoidine biosynthesis, ATP and L-glutamine were directly added to the reaction solution to a final concentration of 0.65 mM each, and product formation was followed by spectroscopic analysis. *In vitro* produced BpsA showed slightly slower rates than BpsA recombinantly produced in *E. coli* (diluted in the PURExpress reaction solution). An accelerated reaction compared to the IVPS conditions was observed for recombinant BpsA in assay buffer, implying that components in the reaction solution compromise catalytic activity. (Figure 3A & B and Figure S4A & B). We note that the highly homologous NRPS IndC (146.5 kDa) from *Photorhabdus luminescens* was also expressed and phosphopantetheinylated *in vitro* to biosynthesize indigoidine ^29^. However, a quantitative analysis was performed with BpsA only (Figure S4C).

**Figure 3.**
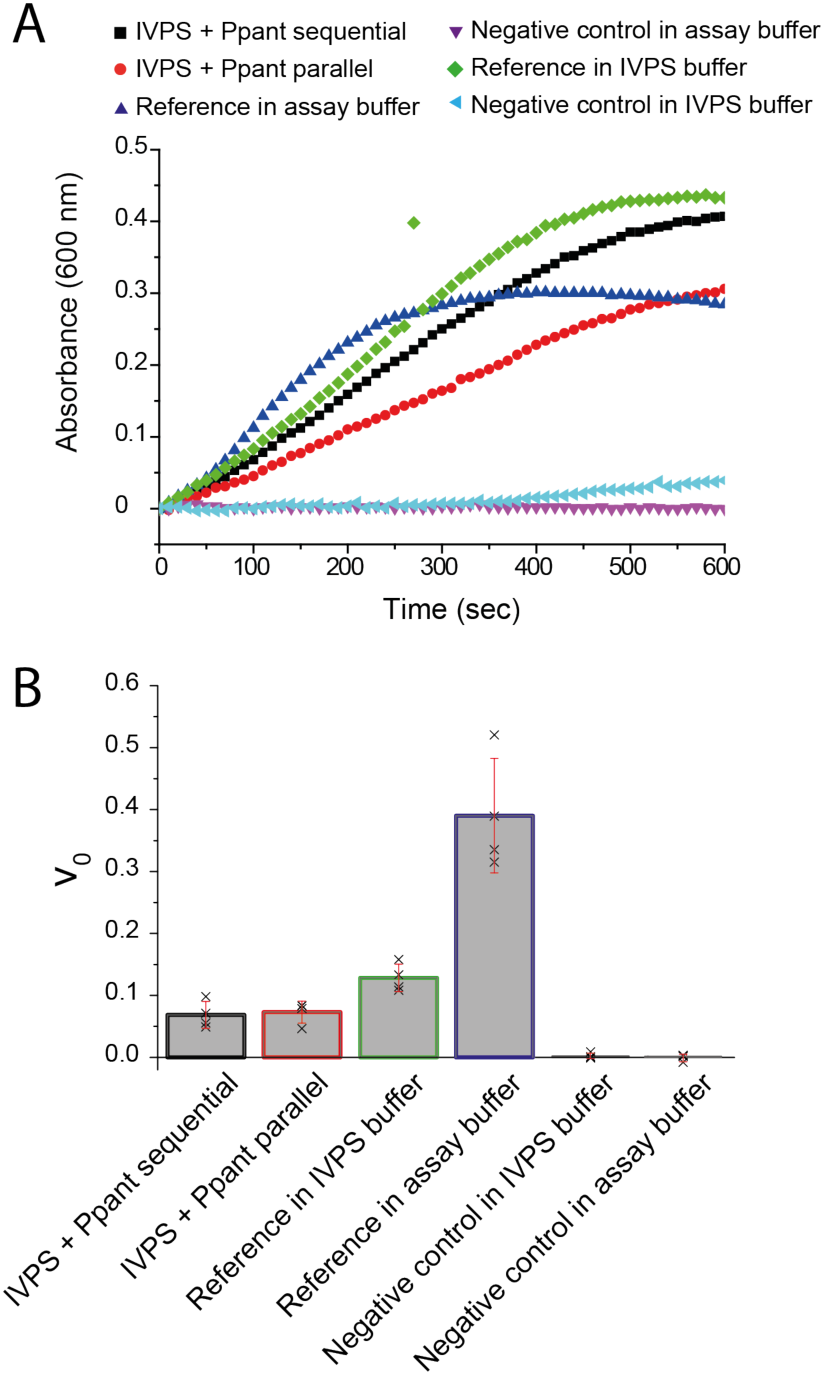
Cell-free synthesis of indigoidine. **(A)** Production of indigoidine spectroscopically followed at 600 nm and compared to turnover rates of each protein preparation (see legend). Data shows one of four independent experiments (for more data, see Figure S4A). **(B)** Turnover rates were calculated taking the maximal slope of product formation rates (curves shown in panel A). Data has been collected on four independent protein preparations. The error bars represent standard deviation (*n* = 4) of maximal rates.

In the light of the successful proof of concept, a set of megasynthases (NRPSs, PKSs, FASs) was screened for synthesis by the PURE cell-free system with coupled phosphopantetheinylation (sequential protocol, see Figure S1). We were able to synthesize and CoA-647-label all megasynthases applied in this screen, *Penicillium patulum* MSAS ^30^, RAPS module 14 ^31^, PikA module 5 (PikAIII) ^32^, DEBS module 4 ^7^, three variants of murine FAS (wild-type, KS-MAT-ACP-TE and with C-terminal GFP (FAS-GFP)) ^33-34^, GrsA ^35^, TycB1 ^36^, and the *Xhenorhabdus* KJ12.1 KJ12ABC ^21^ (Figure S5A & B). The murine FAS construct DH-KR-TE and the NRPS module KJ12C, which do not harbor a carrier protein domain, were not fluorescently labeled, as expected (Figure S5B). The successful phosphopantetheinylation suggests that proteins are correctly folded.

In seeking to probe the PURE cell-free system for the production of another megasynthase, we decided to work with the RXP-synthesizing NRPS KJ12ABC from the bacterium *Xhenorhabdus* KJ12.1^21^. The RXP biosynthetic gene cluster of *Xhenorhabdus* KJ12.1 encodes three NRPS modules – *kj12A* encoding a C-A-T module (137.8 kDa), *kj12B* encoding a C-A/MT-T module (181.5 kDa), and *kj12C* encoding a stand-alone C_term_ domain (62.3 kDa) (see Figure 1B). The RXP-synthesizing NRPS represented the most complex megasynthase system used in our study, and the individual proteins were available in good yields as identified in our screen (see Figure S5B). As a reference for the analysis of KJ12ABC form the PURE cell-free system, modules were recombinantly expressed in *E. coli* and subjected to a product synthesis assay (Figure S6). Specifically, we focused on KJ12B, which was available in second highest yields after the indigoidine-producing monomodular BpsA and IndC (see Figure S5B). The production yield of modules KJ12B and C were estimated from densitometric analysis. The degree of phosphopantetheinylation for KJ12B was determined with the fluorescent analogue CoA-647 and in-gel fluorescence intensities, similarly as performed with BpsA. In order to avoid a compromised activity of the *in vitro* synthesized proteins in the PURExpress reaction solution, as observed for BpsA, we reverse-purified KJ12B and C by Ni-chelating chromatography. Since all proteins from the PURE system (except for the ribosomes) are His-tagged, the Strep-tagged *in vitro* synthesized NRPS modules appear in the flow through, allowing for rapid purifications (Figure 4A). Module KJ12A was added in excess as recombinantly produced and purified protein. We expected that elevated concentrations of the first, chain-initiating module KJ12A shifts the output spectrum to shorter peptides (with less methylated valine residues) at high abundance. Under the reaction conditions, which corresponds to a molar ratio of 10:1:1 (KJ12A:B:C) for the modules, we eventually identified three peptides; the tetrapeptide mV-V-mV-mV-PEA (RXP number 3, see Figure S6) in highest abundance and also two other tetrapeptides of sequences V-V-V-mV-PEA (9) and V-mV-V-mV-PEA (10), respectively (Figure 4B & Figure S7). Since KJ12A alone and in combination with KJ12C cannot produce those peptides (see Figure S6) our data reveal the functionality of the *in vitro* synthesized components KJ12B and C, and underline the suitability of the PURE system as IVPS platform for megasynthase production and analysis.

**Figure 4.**
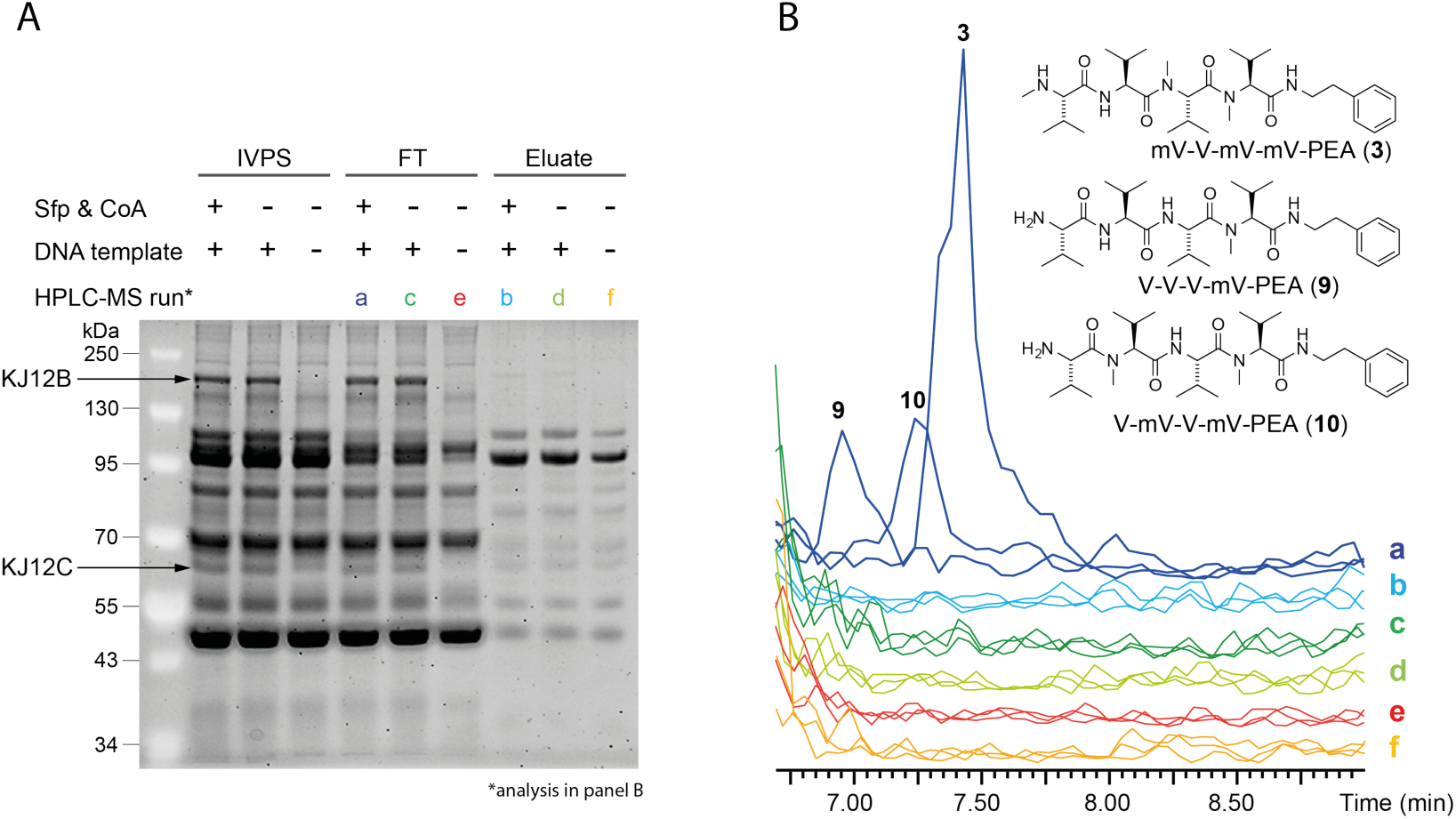
Cell-free synthesis of rhabdopeptides. **(A)** SDS-PAGE of IVPS of KJ12B and KJ12C. Composition of reaction solutions as indicated. **(B)** Specific extracted-ion chromatograms (EICs) of the different HPLC-MS analyses color coded as outlined in (A). The structures of detected rhabdopeptides are attached. For MS-MS fragmentation data of compounds 3, 9 and 10, see Figure S7.

In conclusion, we demonstrate that the PURE system can provide a superior setting for the analysis and engineering of biosynthetic pathways, which is not possible with cell-free synthesis in *E. coli* cell extract ^20^. We show that IVPS can pave the way towards a rapid cell-free screening platform for natural compounds discovery: Genome mining identifies the (megasynthase) gene cluster → gene synthesis provides the DNA → the proteins are produced in the PURE system → enzymatic properties and product output are directly monitored. In spite of the advantages of IVPS by the PURE system, its broad application hinges on solutions to several key challenges. Providing the protein in sufficient yields and quality may be the biggest current limitation: In probing the synthesis of several megasynthases, we observe different yields (see Figure S5A & B), and the in-depth analysis of *in vitro* BpsA production indicates that the quality of the protein is slightly compromised (see Figure 3A & B). Yields and quality of proteins may further be improved by optimizing the encoding DNA in untranslated and translated regions, e.g., in the secondary structure of DNA/RNA or in codon usage ^37-38^. A broader reconstitution of the protein quality control machinery is generally possible, but will need to be balanced against the benefit of a low complex environment for controlling and monitoring reaction progress and output. In this respect, the capability of *E. coli* to produce an array of megasynthases is good news for the applicability of the *E. coli* based PURE system ^6^. It implies that the folding and assembly of megasynthases do generally not rely on specific factors of original producer strains. A further limitation arises from the current high costs for IVPS, particularly of systems reconstituted from pure components. However, recent work promises that the access to PURE cell-free systems will improve and become less costly in the near future ^39^.

## Methods

### Recombinant production of of BpsA *in vivo*

The *bpsA* gene with N-term Strep-tag was placed in a pET22b plasmid with amp resistance. BpsA was expressed in BL21 Gold (DE3) *E. coli* cells, in TB medium at 20 °C, 180 rpm overnight. Cells were disrupted mechanically using a French press at 1000 bar. Strep-tagged protein was purified over Strep-Tactin Sepharose columns (iba) in Strep-buffer (20 mM Tris-HCl, 500 mM NaCl, 1 mM EDTA, pH7.9), eluted with 2.5 mM D-desthiobiotin. Purified BpsA was dialyzed against assay buffer (50 mM HEPES-KOH, 100 mM monopotassium L-glutamate, 13 mM magnesium acetate, pH7.6) overnight. Size exclusion chromatography was performed on Superdex 200 gel filtration columns. The protein was concentrated in Amicon Ultra centrifugal filters, MWCO 10k (Merck Millipore). For DNA sequence encoding BpsA and IndC, see Figure S8.

### Recombinant production of KJ12A, KJ12B and KJ12C

For production of KJ12A, KJ12B and KJ12C, overnight cultures from *E. coli* BL21 (DE3) Star™mtaA strains containing pCOLA_KJ12A_His/Strep and pCOLA_KJ12B_Strep as well as *E. coli* BL21 (DE3) Star™containing pCDF_KJ12C_Strep were separately transferred into 1 L fresh LB medium supplemented with kanamycin, chloramphenicol (KJ12A and KJ12B) and spectinomycin (KJ12C), respectively. The medium was mixed with 50 mL 20×NPS (66 g (NH_4_)_2_SO_4_, 136 g H_2_PO_4_, 142 g Na_2_HPO_4_ and 900 mL ddH_2_O, sterile filtrated), 20 mL 50×5052 (250 g glycerol, 25 g glucose, 100 g α-Lactose and 730 mL ddH_2_O, sterile filtrated), 1 mL MgSO_4_ for auto induction ^40^ and 5 mL 10 % L-arabinose for induction of the PPtase (MtaA) production (KJ12A and KJ12B), followed by incubation at 37 °C with shaking at 180 rpm. After the cultures grew to an OD_600_ of 0.5-0.7, they were kept growing for 72 h at 180 rpm and 20 °C. The cells were harvested by centrifugation (10,000 rpm, 10 min, 4 °C). For purification, cell pellets were resuspended in 50–100 mL Strep-tag binding buffer (100 mM Tris-HCl, 150 mM NaCl, pH 8.0, sterile filtrated) supplemented with one protease inhibitor tablet (Roche), 0.1 % of Triton X-100, 0.5 mg/mL lysozyme (10 U/mL) und 3 μL Benzonase Nuklease^®^ (25 U/μL) and incubated for 30 min at rt and additionally lysed by sonication. Cell debris was removed from the lysate by centrifugation (20,000 rpm, 30 min, 4 °C) and the lysate was loaded onto a StrepTrap HP 5 mL column (GE Healthcare) and purified with the ÄKTA™purifier system (GE Healthcare) or NGC system (BioRad). 50–100 mL protein lysate was loaded onto the column, which was equilibrated with two column volumes (CV) of Strep-tag binding buffer (100 mM Tris-HCl, 150 mM NaCl, pH 8.0, sterile filtrated), with a flow rate of 2.5 mL/min and fraction collection of 12 mL. The column was washed with 12 CV Strep-tag binding buffer with a flow rate of 5 mL/min to wash away unspecifically bound proteins. Protein of interest was eluted with 8 CV of Strep-tag elution buffer (100 mM Tris-HCl, 150 mM NaCl, 2.5 mM desthiobiotin, pH 8.0, sterile filtrated) with a flow rate of 5 mL/min and fractions of 5 mL were collected. The column was regenerated with 3 CV regeneration buffer (Strep-tag binding buffer with 1 mM HABA, pH 8.0; sterile filtrated) and additional 5 CV Strep-tag binding buffer. The pooled fractions containing the protein of interest were pooled and concentrated with Centriprep units, MWCO= 50 kDa (Merck Millipore) for KJ12A and KJ12B or MWCO = 30 kDa (Merck Millipore) for KJ12C.

### *In vitro* protein synthesis and phosphopantetheinylation

IVPS was performed using PURExpress *In Vitro* Protein Synthesis Kit (NEB) with supplements (16 U RNase inhibitor (NEB), 5 mM FMN). 50-100 ng DNA template was provided in 25 µL reaction volume and incubated for 2 h at 30 °C. The reaction was stopped on ice, adding 50 µg/mL kanamycin as ribosome inhibitor. The IVPS reaction could also be scaled up to 100 µL. Phosphopantetheinylation was either performed in parallel to the IVPS or after IVPS by adding Sfp (final concentration 0.5 µM) and CoA (final concentration 100 µM) to the reaction. For simultaneous phosphopantetheinylation (parallel protocol) the reaction mixture was incubated another 15 min at 30 °C after ribosome inhibition by kanamycin (final concentration of 50 µg/ml, for subsequent phosphopantetheinylation for 1 h at 30 °C. The reaction was stopped on ice. Phosphopantetheinylation was quantified from in-gel fluorescence intensities, using fluorescent CoA 647 (NEB) as substrate. Protein expression yields were quantified from SDS-PAGE by densitometric analysis using unfolded *in vivo* expressed and purified apo-BpsA as reference. Western blots were treated with antibodies against the Strep-tag and analyzed by fluorescence detection. As primary antibody StrepMAB-Classic from mouse (IBA), and as secondary antibody Donkey anti-Mouse IgG DyLight 755 conjugate (Thermo Fisher Scientific) were used.

### Indigoidine synthesis

Cell-free synthesis of indigoidine synthesis was performed by adding 0.65 mM ATP and L-glutamine and 0.8 mM MgCl_2_ to the phosphopantetheinylated IVPS mixture and incubating for 1 h at RT. The synthesis of indigoidine was monitored measuring the absorbance at 600 nm over time, in cuvettes at the NanoDrop (Thermo Fisher Scientific) or in 384 well plates at the CLARIOstar (BMG). Turnover rates were determined by calculating the point of maximal slope (inflection point) of the sigmoidal curves by the second derivation and normalized to protein concentration.

### RXP synthesis

For *in vitro* reactions, 10 mM MgCl_2_, 5 mM L-valine, 2 mM *S*-adenosyl-methionine (SAM) and 3 mM PEA were first added in 1.5 mL reaction tubes. The proteins KJ12A, KJ12B and KJ12C with ratios of KJ12A:KJ12C 1:1 (2 μM/2 μM), KJ12A:KJ12B 1:1 (2 μM/2 μM), KJ12B:KJ12C 1:10 (2 μM/20 μM), 1:1 (2 μM/2 μM), and 10:1 (20 μM/2 μM), as well as KJ12A:KJ12B:KJ12C 1:1:1 (2 μM/2 μM/2 μM/) and 1:10:1 (2 μM/20 μM/2 μM/) were then separately added into the above mentioned reaction tubes to check the RXP profiles. Finally, the total volume was adjusted to 400 µL with assay buffer (50 mM Tris-HCl, 50 mM NaCl, pH 8.0) and the reaction was started by addition of 3 mM ATP. The negative controls, one without all enzymes and the other with KJ12A, KJ12B or KJ12C alone were performed in the same way. All reaction tubes were incubated at 22.5 °C, 250 rpm for 20 h. After incubation, the samples were extracted with one volume of methanol for 1 h at rt under shaking and measured with the HPLC-MS analysis. Cell-free synthesis was performed in reaction volumes of 125 µL. Phosphopantetheinylation was performed in sequential manner. By taking a PUREexpress protein band as reference, determined in concentration before with the BpsA calibration curve, the concentrations of KJ12B and KJ12C were determined to 0.02 and 0.015 µM, respectively. After cell-free synthesis, reaction solutions for modules KJ12B and KJ12C were combined and inversely purified with Ni-NTA Magnetic Beads (Thermo Fisher) using assay buffer containing 30 mM imidazole and 0.05 % Tween-20. Strep-tagged NRPS modules were collected in the flow through. Elution of His-tagged proteins from the reaction solution was performed with assay buffer containing 200 mM Imidazole and 0.05 % Tween-20. The elution fraction served as negative control. For cell-free synthesis, recombinantly produced KJ12A was added to the inversely purified combined reaction solution of KJ12B and C at a final concentration of 0.16 µM. Accordingly, the molar ratio of modules in the reaction solution was approximately 10:1:1 (A:B:C). The reaction was developed and prepared for MS-analysis as described below.

### HPLC-MS analysis of RXP production

5 μl of the crude extracts were injected and analyzed via ESI-HPLC-MS by a Dionex UltiMate 3000 HPLC system coupled to a Bruker AmaZon X mass spectrometer with a ACQUITY UPLC™ BEH C18 column (130 Å, 2.1 mm × 100 mm, 1.7 μm particle size, Waters GmbH) at a flow rate of 0.6 mL/min for 16 min, using acetonitrile and water supplemented with 0.1 % formic acid (v/v) in a gradient ranging from 5 % to 95 % of acetonitrile (ACN). For RXPs detection positive mode with scanning range from 100-1200 *m/z* and UV at 200-600 nm was used. The software DataAnalysis 4.3 (Bruker) was used to evaluate the generated HPLC-MS measurements.

## Acknowledgments

We thank Ines Gössner for technical assistance in this project.

## Authors contribution

I.S. performed experiment to IVPS, analyzed data, and reviewed and edited the manuscript, S.N. purified KJ12A/B/C and performed experiment to *in vitro* production of rhabdopeptides, analyzed data, and wrote the manuscript, C.H. performed experiment to IVPS, analyzed data, and wrote the manuscript, N.C. performed initial experiments, P.T. performed experiments, and conceived the project, H.B.B. designed research to RXP-producing NRPS KJ12ABC, analyzed data, and reviewed and edited the manuscript, M.G. designed research to cell-free synthesis of natural compounds, analyzed data, and wrote the manuscript.

## Table of Contents

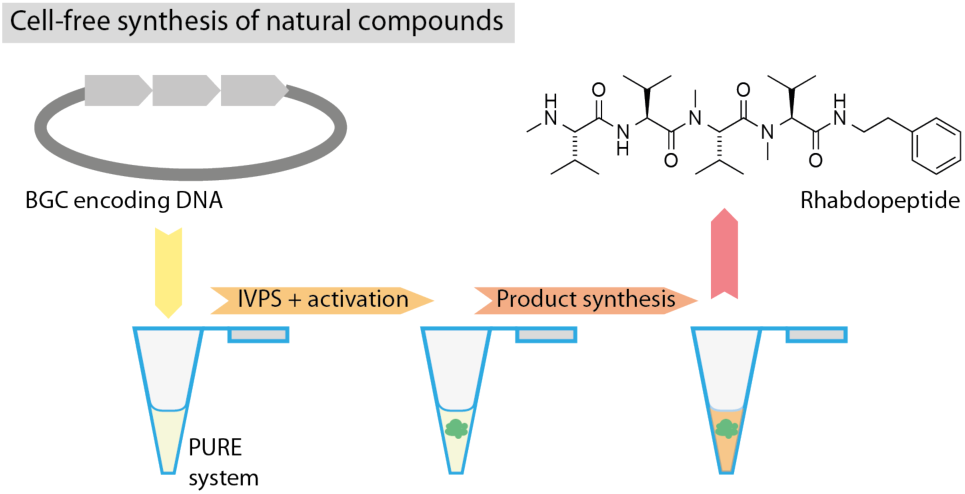

## References

1. Ziemert, N., Alanjary, M., and Weber, T. (2016) The evolution of genome mining in microbes - a review. Nat. Prod. Rep. 33, 988–1005.

2. Dejong, C. A., Chen, G. M., Li, H., Johnston, C. W., Edwards, M. R., Rees, P. N., Skinnider, M. A., Webster, A. L., and Magarvey, N. A. (2016) Polyketide and nonribosomal peptide retro-biosynthesis and global gene cluster matching. Nat. Chem. Biol. 12, 1007–1014.

3. Stewart, E. J. (2012) Growing Unculturable Bacteria. J. Bacteriol. 194, 4151–4160.

4. Zhang, M. M., Wang, Y., Ang, E. L., and Zhao, H. (2016) Engineering microbial hosts for production of bacterial natural products. Nat. Prod. Rep. 33, 963–987.

5. Huo, L., Hug, J. J., Fu, C., Bian, X., Zhang, Y., and Muller, R. (2019) Heterologous expression of bacterial natural product biosynthetic pathways. Nat. Prod. Rep. 36, 1412–1436.

6. Shimizu, Y., Inoue, A., Tomari, Y., Suzuki, T., Yokogawa, T., Nishikawa, K., and Ueda, T. (2001) Cell-free translation reconstituted with purified components. Nat. Biotechnol. 19, 751–755.

7. Pieper, R., Luo, G., Cane, D. E., and Khosla, C. (1995) Cell-free synthesis of polyketides by recombinant erythromycin polyketide synthases. Nature 378, 263–266.

8. Ku, J., Mirmira, R. G., Liu, L., and Santi, D. V. (1997) Expression of a functional non-ribosomal peptide synthetase module in Escherichia coli by coexpression with a phosphopantetheinyl transferase. Chem. Biol. 4, 203–207.

9. Fischbach, M. A. and Walsh, C. T. (2006) Assembly-line enzymology for polyketide and nonribosomal Peptide antibiotics: logic, machinery, and mechanisms. Chem. Rev. 106, 3468–3496.

10. Hertweck, C. (2009) The Biosynthetic Logic of Polyketide Diversity. Angew. Chem. Int. Ed. 48, 4688–4716.

11. Sussmuth, R. D., and Mainz, A. (2017) Nonribosomal Peptide Synthesis-Principles and Prospects. Angew. Chem. Int. Ed. Engl. 56, 3770–3821.

12. Cortes, J., Haydock, S. F., Roberts, G. A., Bevitt, D. J., and Leadlay, P. F. (1990) An unusually large multifunctional polypeptide in the erythromycin-producing polyketide synthase of Saccharopolyspora erythraea. Nature 348, 176–178.

13. Weckermann, R., Furbass, R., and Marahiel, M. A. (1988) Complete nucleotide sequence of the tycA gene coding the tyrocidine synthetase 1 from Bacillus brevis. Nucleic Acids Res. 16, 11841.

14. Khosla, C., (1997) Harnessing the Biosynthetic Potential of Modular Polyketide Synthases. Chem. Rev. 97, 2577–2590.

15. Marahiel, M. A., Stachelhaus, T., and Mootz, H. D. (1997) Modular Peptide Synthetases Involved in Nonribosomal Peptide Synthesis. Chem. Rev. 97, 2651–2674.

16. Klaus, M., and Grininger, M. (2018) Engineering strategies for rational polyketide synthase design. Nat. Prod. Rep. 35, 1070–1081.

17. Yuzawa, S., Backman, T. W. H., Keasling, J. D., and Katz, L. (2018) Synthetic biology of polyketide synthases. J. Ind. Microbiol. Biotechnol. 45, 621–633.

18. Bozhuyuk, K. A., Micklefield, J., and Wilkinson, B. (2019) Engineering enzymatic assembly lines to produce new antibiotics. Curr. Opin. Microbiol. 51, 88–96.

19. Takahashi, H., Kumagai, T., Kitani, K., Mori, M., Matoba, Y., and Sugiyama, M. (2007) Cloning and characterization of a Streptomyces single module type non-ribosomal peptide synthetase catalyzing a blue pigment synthesis. J. Biol. Chem. 282, 9073–9081.

20. Goering, A. W., Li, J., McClure, R. A., Thomson, R. J., Jewett, M. C., and Kelleher, N. L. (2017) In Vitro Reconstruction of Nonribosomal Peptide Biosynthesis Directly from DNA Using Cell-Free Protein Synthesis. ACS Synth. Biol. 6, 39–44.

21. Hacker, C., Cai, X., Kegler, C., Zhao, L., Weickhmann, A. K., Wurm, J. P., Bode, H. B., and Wöhnert, J. (2018) Structure-based redesign of docking domain interactions modulates the product spectrum of a rhabdopeptide-synthesizing NRPS. Nat. Commun. 9, 4366.

22. Reverchon, S., Rouanet, C., Expert, D., and Nasser, W. (2002) Characterization of indigoidine biosynthetic genes in Erwinia chrysanthemi and role of this blue pigment in pathogenicity. J. Bacteriol. 184, 654–665.

23. Schmidt, T., and Skerra, A. (2007) The Strep-tag system for one-step purification and high-affinity detection or capturing of proteins. Nat. Protoc. 2, 1528–1535.

24. Walsh, C. T., Gehring, A. M., Weinreb, P. H., Quadri, L. E., and Flugel, R. S. (1997) Post-translational modification of polyketide and nonribosomal peptide synthases. Curr. Opin. Chem. Biol. 1, 309–315.

25. Quadri, L. E., Weinreb, P. H., Lei, M., Nakano, M. M., Zuber, P., and Walsh, C. T. (1998) Characterization of Sfp, a Bacillus subtilis phosphopantetheinyl transferase for peptidyl carrier protein domains in peptide synthetases. Biochemistry 37, 1585–1595.

26. Lee, K. K., Da Silva, N. A., and Kealey, J. T. (2009) Determination of the extent of phosphopantetheinylation of polyketide synthases expressed in Escherichia coli and Saccharomyces cerevisiae. Anal. Biochem. 394, 75–80.

27. Beld, J., Sonnenschein, E. C., Vickery, C. R., Noel, J. P., and Burkart, M. D. (2014) The phosphopantetheinyl transferases: catalysis of a post-translational modification crucial for life. Nat. Prod. Rep. 31, 61–108.

28. Worthington, A. S., and Burkart, M. D. (2006) One-pot chemo-enzymatic synthesis of reporter-modified proteins. Org. Biomol. Chem. 4, 44–46.

29. Brachmann, A. O., Kirchner, F., Kegler, C., Kinski, S. C., Schmitt, I. and Bode, H. B., (2012) Triggering the production of the cryptic blue pigment indigoidine from Photorhabdus luminescens. J. Biotechnol. 157, 96–99.

30. Dimroth, P., Ringelmann, E., and Lynen, F. (1976) 6-Methylsalicylic acid synthetase from Penicillium patulum. Some catalytic properties of the enzyme and its relation to fatty acid synthetase. Eur. J. Biochem. 68, 591–596.

31. Schwecke, T., Aparicio, J. F., Molnár, I., König, A., Khaw, L. E., Haydock, S. F., Oliynyk, M., Caffrey, P., Cortés, J., Lester, J. B., Böhm, G. A., Staunton, J., and Leadlay, P. F. (1995) The biosynthetic gene cluster for the polyketide immunosuppressant apamycin. Proc. Natl. Acad. Sci. U S A. 92, 7839–7843.

32. Aldrich, C., Beck, B., Fecik, R., and Sherman, D. (2005) Biochemical investigation of pikromycin biosynthesis employing native penta- and hexaketide chain elongation intermediates. J. Am. Chem. Soc. 127, 8441–8452.

33. Rittner, A., Paithankar, K. S., Huu, K. V., and Grininger, M. (2018) Characterization of the Polyspecific Transferase of Murine Type I Fatty Acid Synthase (FAS) and Implications for Polyketide Synthase (PKS) Engineering. ACS Chem. Biol. 13, 723–732.

34. Rittner, A., Paithankar, K. S., Drexler, D. J., Himmler, A., and Grininger, M. (2019) Probing the modularity of megasynthases by rational engineering of a fatty acid synthase Type I. Protein Sci. 28, 414–428.

35. Krätzschmar, J., Krause, M., and Marahiel, M. A. (1989) Gramicidin S biosynthesis operon containing the structural genes grsA and grsB has an open reading frame encoding a protein homologous to fatty acid thioesterases. J. Bacteriol. 171, 5422–5429.

36. Mootz, H. D., and Marahiel, M. A. (1997) The tyrocidine biosynthesis operon of Bacillus brevis: complete nucleotide sequence and biochemical characterization of functional internal adenylation domains. J. Bacteriol. 179, 6843–6850.

37. Boel, G., Letso, R., Neely, H., Price, W. N., Wong, K. H., Su, M., Luff, J. D., Valecha, M., Everett, J. K., Acton, T. B., Xiao, R., Montelione, G. T., Aalberts, D. P., and Hunt, J. F. (2016) Codon influence on protein expression in E. coli correlates with mRNA levels. Nature 529, 358–363.

38. Plotkin, J. B., and Kudla, G. (2011) Synonymous but not the same: the causes and consequences of codon bias. Nat. Rev. Genet. 12, 32–42.

39. Lavickova, B., and Maerkl, S. J. (2019) A Simple, Robust, and Low-Cost Method To Produce the PURE Cell-Free System. ACS Synth. Biol. 8, 455–462.

40. Studier, F. W. (2005) Protein production by auto-induction in high density shaking cultures. Protein Expr. Purif. 41, 207–234.

