## Supplemental Material for "Cell-free synthesis of natural compounds from genomic DNA of biosynthetic gene clusters"

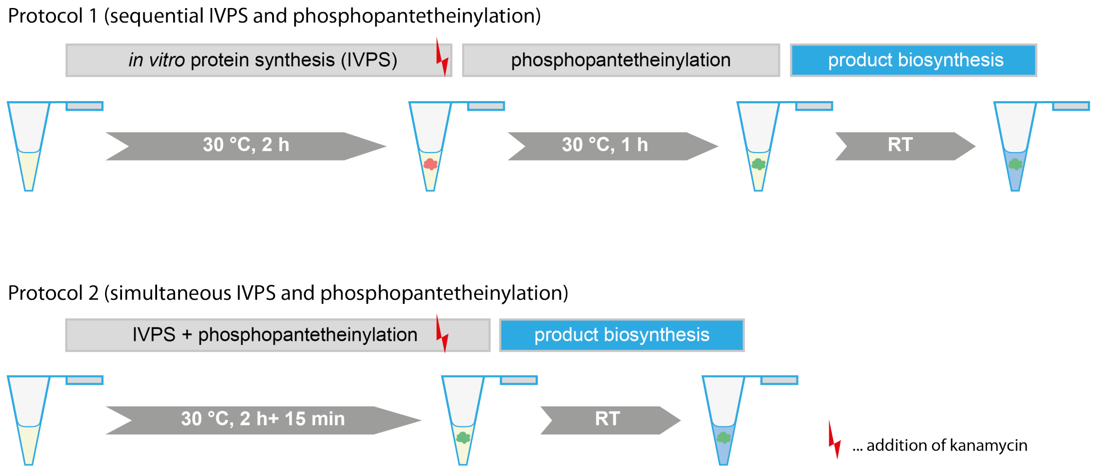


**Figure S1. IVPS protocols used in this study.** Protocol 1: IVPS is performed first and phosphopantetheinylation started after ribosome inhibition by adding CoA and Sfp. Protocol 2: CoA and Sfp are added directly to the cell-free system for simultaneous phosphopantetheinylation and IVPS. The reaction is incubated for another 15 minutes at 30 °C after ribosome inhibition to ensure phosphopantetheinylation of newly synthesized protein.

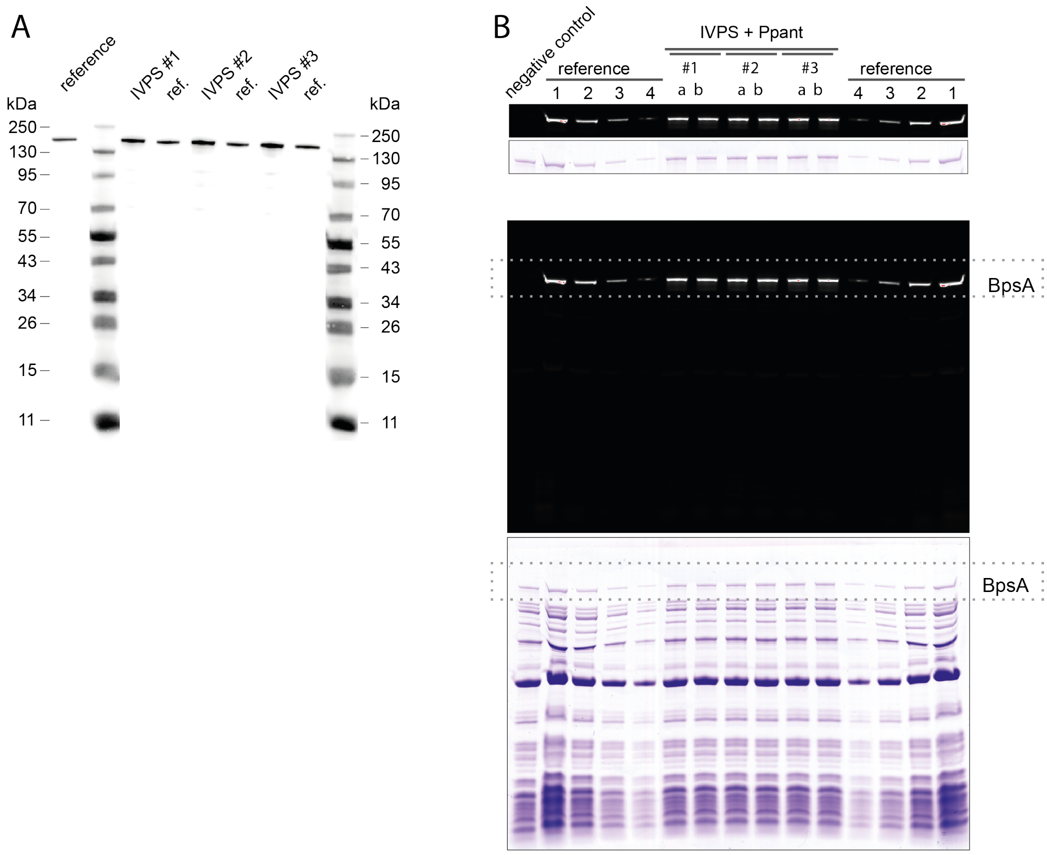


**Figure S2. IVPS of BspA. (A)** Uncropped blot shown in Figure 2A for the expression control by Western blotting with anti-Strep antibodies performed in three independent reaction solutions (#1-3). BspA was applied as *holo*-protein; i. e., first produced by IVPS with simultaneous phosphopantetheinylation by the addition of *B. subtilis* Sfp. Self-cast 9 % Tris-Tricine gel. **(B)** Uncropped gels shown in Figure 2C for the quantification of protein production yields and phosphopantetheinylation efficiency. BpsA was first produced by IVPS and then phosphopantetheinylated with Sfp and CoA-647 (purchased from NEB). Samples from three independent reactions (#1-3) were applied in repetition (a & b). For calibration, recombinantly produced BpsA, diluted in the IVPS reaction solution, was loaded in amounts of 1.25, 0.625, 0.3125 and 0.15625 pmol. 9 % Tris-Tricine gel as in panel A. Areas of the gel shown in Figure 2C are highlighted with dashed lines.

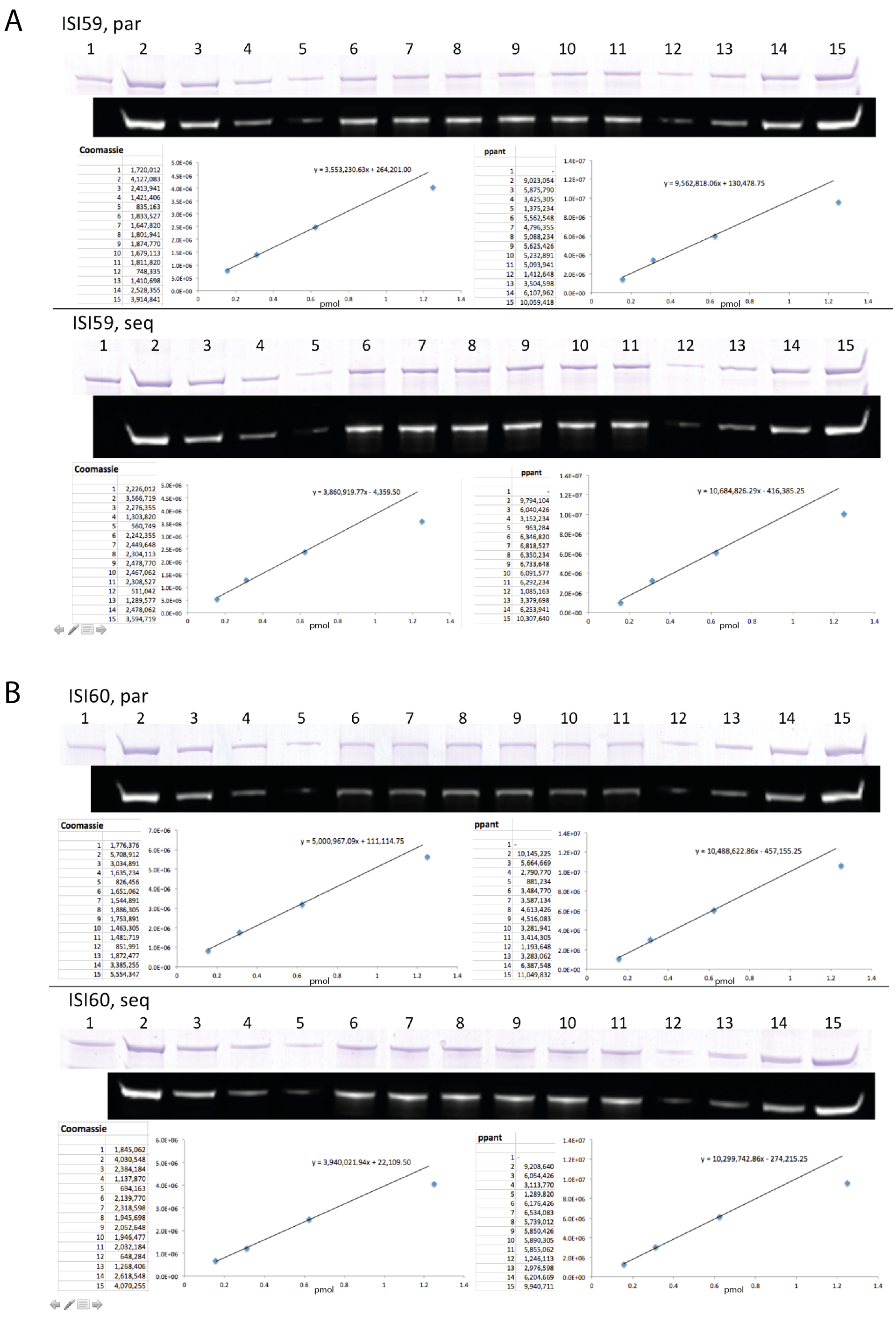

A
ISI59, par
y = 3,553,230.63x + 264,201.00
y = 9,562,818.06x + 130,478.75
ISI59, seq
y = 3,860,919.77x - 4,359.50
y = 10,684,826.29x - 416,385.25
B
ISI60, par
y = 5,000,967.09x + 111,114.75
y = 10,488,622.86x - 457,155.25
ISI60, seq
y = 3,940,021.94x + 22,109.50
y = 10,299,742.86x - 274,215.25
Coomassie
ppant
pmol


**(Figure S3)**

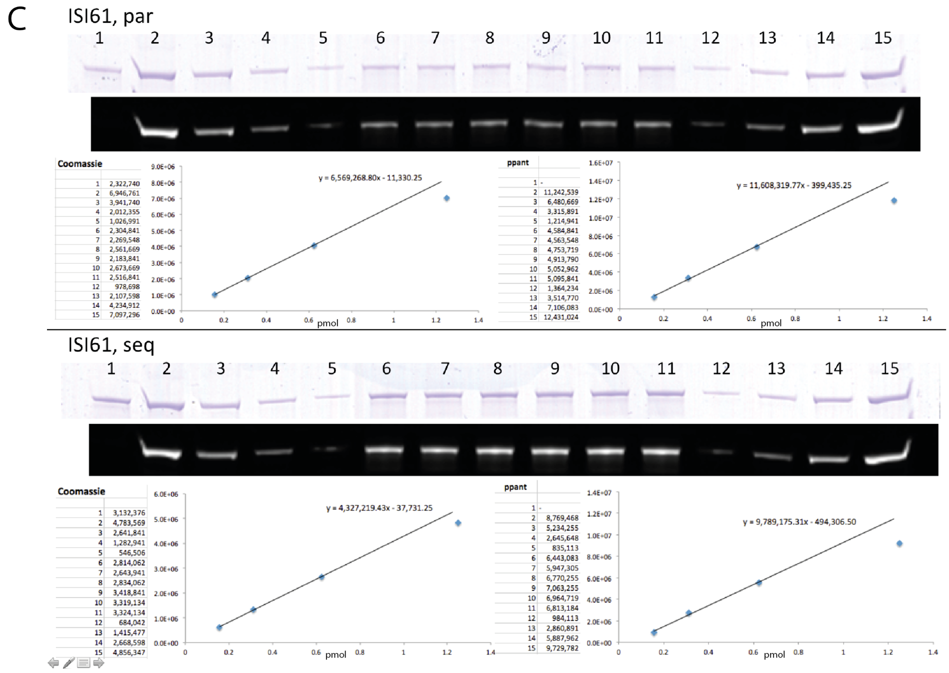


**Figure S3 (continued). Gels for quantification.** Gel of three independent reactions **(A, B and C)** with each reaction conducted in duplicate. Labeling (ISI59-61) refer to internal experiment numbers. Lanes: 1, recombinantly produced BpsA; lanes 2-5 and 12-15, BpsA diluted in the IVPS reaction solution and loaded at amounts of 1.25 (2 and 15), 0.625 (3 and 14), 0.3125 (4 and 13) and 0.15625 pmol (5 and 12); 6 and 7, IVPS reaction #1; 8 and 9, IVPS reaction #2; 10 and 11, IVPS reaction #3. Reactions were performed with the parallel protocol (termed “par”) and the sequential protocol (“seq”). 9 % Tris-Tricine gel were used. Phosphopantetheinylation was performed with Sfp and CoA-647 (purchased from NEB).

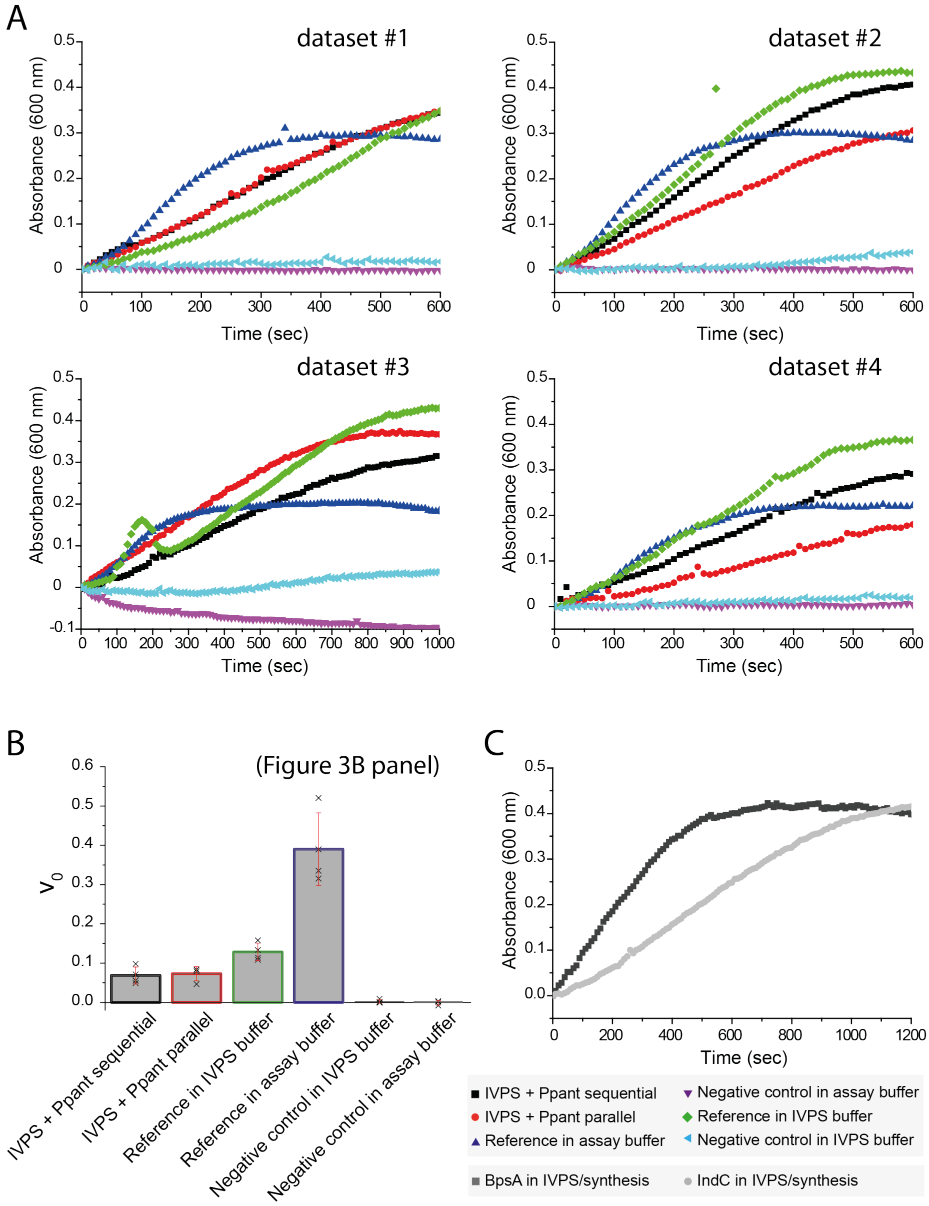


**Figure S4. Synthesis of indigoidine. (A)** Extended data to Figure 3A. Production of indigoidine was spectroscopically followed at 600 nm to compare turnover rates of each protein preparation (see legend). Production was followed over a time range of 1 h. The drop in OD may originate from solubility problems or effects caused by the oxidative maturation of the compound. **(B)** Turnover rates were calculated using linear regression. Identical to panel Figure 3B, shown here for clarity. **(C)** Qualitative test of IndC activity after IVPS (compared to BpsA).

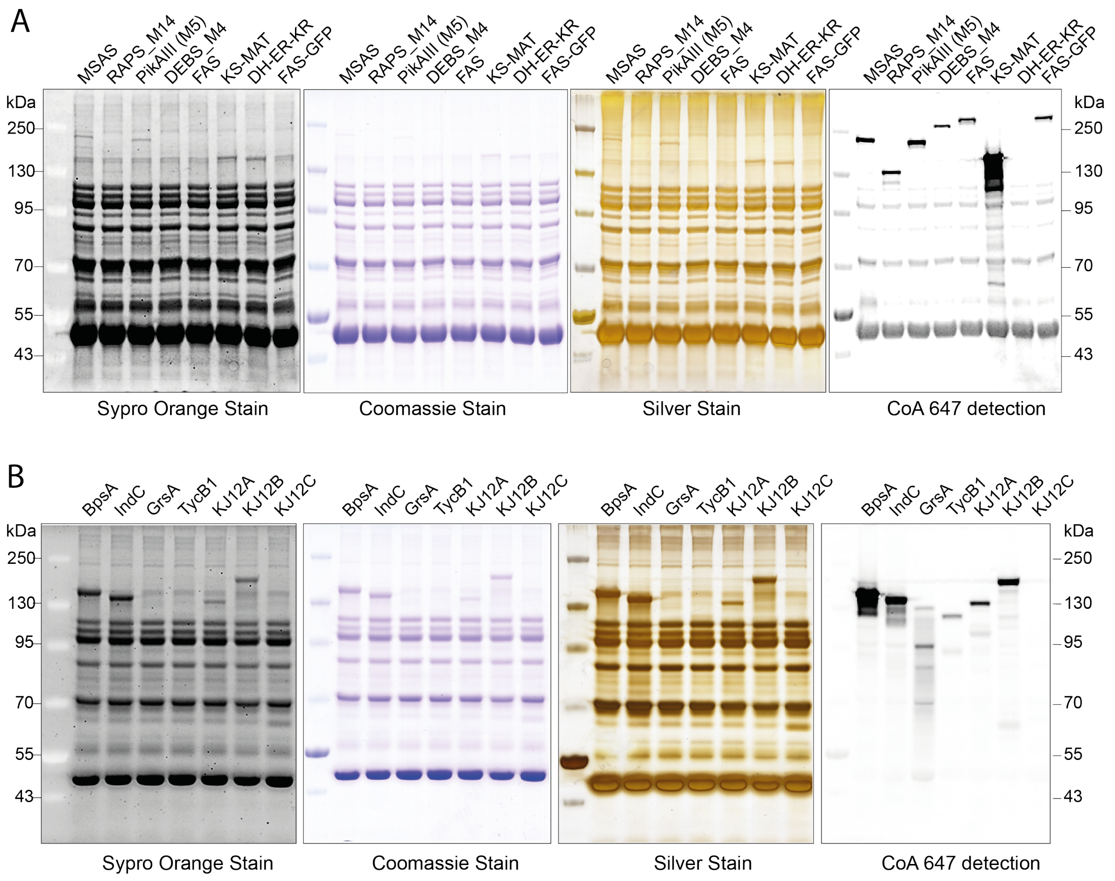


**Figure S5. IVPS screen of selected megasynthases. (A)** IVPS and phosphopantetheinylation of PKSs and mouse FAS (A) and NRPSs. We note that phosphopantetheinylation of the FAS-construct DH-ER-KR is not possible, as expected, due to the missing ACP. (B). IVPS was performed by the sequential protocol (see Figure S1). SDS-PAGE gels were stained differently. Phosphopantetheinylation were performed with Sfp and CoA-647 (purchased from NEB) and read out by in-gel fluorescence. 9 % Tris-Tricine gels were used. We note that phosphopantetheinylation of the KJ12C is not possible, as expected, due to the missing ACP.

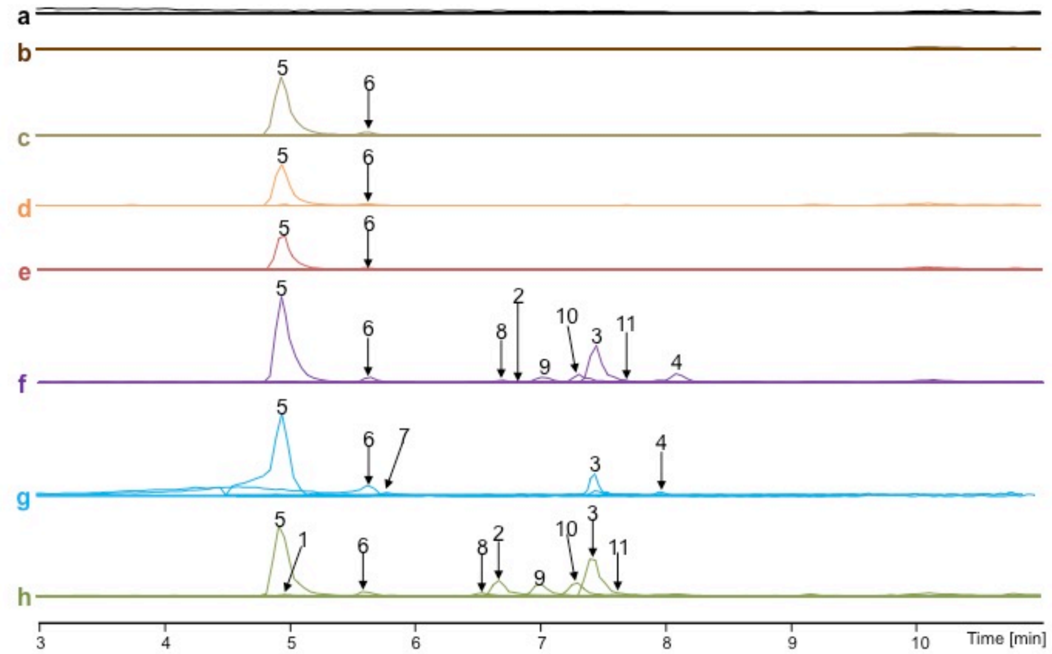


| sample a | b | c | d | e | f | g | h | RXP composition | RXP number |
|---|---|---|---|---|---|---|---|---|---|
| - | - | - | - | - | - | - | ✓ | mV-PEA | 1 |
| - | - | - | - | - | ✓ | - | ✓ | mV-V-mV-PEA | 2 |
| - | - | - | - | - | ✓ | ✓ | ✓ | mV-V-mV-mV-PEA | 3 |
| - | - | - | - | - | ✓ | ✓ | - | mV-V-mV-mV-mV-PEA | 4 |
| - | - | ✓ | ✓ | ✓ | ✓ | ✓ | ✓ | V-PEA | 5 |
| - | - | ✓ | ✓ | ✓ | ✓ | ✓ | ✓ | V-V-PEA | 6 |
| - | - | - | - | - | - | ✓ | - | V-V-V-PEA | 7 |
| - | - | - | - | - | ✓ | - | ✓ | V-V-mV-PEA | 8 |
| - | - | - | - | - | ✓ | - | ✓ | V-V-V-mV-PEA | 9 |
| - | - | - | - | - | ✓ | - | ✓ | V-mV-V-mV-PEA | 10 |
| - | - | - | - | - | ✓ | - | ✓ | V-mV-V-V-mV-PEA | 11 |

**Legend**

**a) Negative control without enzymes**
**b) KJ12C**
**c) KJ12B**
**d) KJ12A**
**e) KJ12A:KJ12C 1:1**
**f) KJ12A:KJ12B 1:1**
**g) KJ12B:KJ12C 1:1**
**h) KJ12A:KJ12B:K12JC 1:1:1**

**Figure S6. Rhabdopeptides production assay with proteins KJ12A, KJ12B and KJ12C from recombinant production in *E. coli*.** (Top) Specific EICs of the different HPLC-MS analyses color coded as outlined in legend. (Bottom) Table of rhabdopeptides produced by recombinantly expressed proteins at various stoichiometry (see legend). For MS-MS fragmentation data, see Cai, X., Nowak, S., Wesche, F. *et al.* [1]

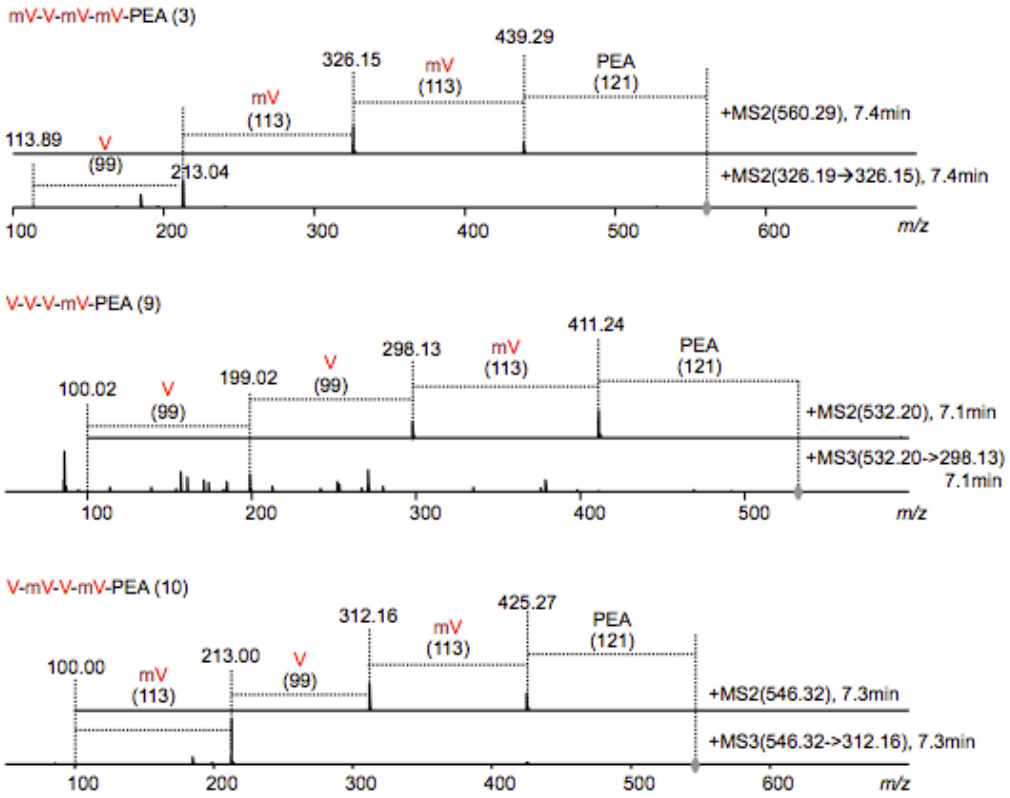


**Figure S7. MS-MS fragmentation of compounds 3, 9 and 10**. This data has been collected from *in vitro* assays with purified proteins.

### BpsA

```
cccgcgaaattaatacgactcactataggggaattgtgagcggataacaattcccctctagaaataattttgtttaac
tttaagaaggagatatacatATGGCTAGCTGGAGCCACCCGCAGTTCGAAAAAGGCGTCGACACTCTTCAGGAGACCA
GCGTGCTCGAGCCCACCCTGCAGGGGACCACCACGTTGCCCGGCCTGCTCGCCCAGCGGGTGGCCGAACACCCCGAGG
CGATCGCGGTCGCCTACCGGGACGACAAGCTCACCTTCCGCGAGCTCGCGTCCAGAAGCGCGGCCCTCGCCGACTACC
TGGAGCACCTCGGTGTCTCCGCCGACGACTGCGTCGGCCTGTTCGTCGAGCCGTCGATCGACCTGATGGTCGGCGCCT
GGGGCATCCTCAACGCCGGCGCCGCGTACCTGCCGCTGTCCCCGGAGTACCCCGAGGACCGGCTGCGCTACATGATCG
AGAACAGCGAGACGAAGATCATCCTGGCGCAGCAGCGCCTGGTGTCCCGTCTGCGCGAGCTCGCGCCGAAGGACGTCA
CCATCGTGACCCTGCGCGAGTCCGAGGCCTTCGTCCGCCCCGAGGGCACCGAGGCCCCGGCCGCCCGCAGCGCCCGCC
CGGACACCCTCGCGTACGTCATCTACACCTCCGGCAGCACGGGCAAGCCGAAGGGTGTGATGATCGAGCACCGCAGCA
TCGTCAACCAGCTCGGCTGGCTGCGCGAGACCTACGCGATCGACCGCAGCAAGGTCATCCTCCAGAAGACCCCGATGA
GCTTCGACGCCGCCCAGTGGGAGATCCTCTCCCCGGCCAACGGCGCCACCGTCGTCATGGGCGCCCCGGGCGTCTACG
CCGACCCCGAGGGCCTCATCGAGACCATCGTCAAGCACAACGTGACCACCCTCCAGTGCGTCCCGACGCTGCTCCAGG
GTCTGATCGACACCGAGAAGTTCCCCGAGTGCGTCTCCCTCCAGCAGATCTTCAGCGGTGGCGAGGCCCTCTCCCGCC
TGCTGGCGATCCAGACCACGCAGGAGATGCCCGGCCGGGCGCTCATCAACGTCTACGGGCCGACCGAGACGACGATCA
ACTCGTCCTCGTTCCCCGTCGACCCCGCCGACCTGGACGAGGGACCGCAGTCCATCTCCATCGGCTCCCCGGTGCACG
GCACCACGTACCACATCCTTGACAAGGAGACCCTCAAGCCGGTCGGCGTCGGTGAGATCGGCGAGCTGTACATCGGCG
GCATCCAGCTGGCCCGCGGCTACCTGCACCGCGACGACCTGACCGCCGAGCGCTTCCTGGAGATCGAGCTCGAGGAGG
GCGCCGAGCCCGTCCGCCTGTACAAGACGGGCGACCTCGGCCAGTGGAACAACGACGGCACCGTGCAGTTCGCCGGCC
GCGCCGACAACCAGGTCAAGCTGCGCGGCTACCGCGTCGAGCTCGACGAGATATCCCTGGCGATCGAGAACCACGACT
GGGTCCGCAACGCCGCCGTCATCGTCAAGAACGACGGCCGCACCGGCTTCCAGAACCTGATCGCCTGCATCGAGCTGA
GCGAGAAGGAAGCCGCCCTGATGGACCAGGGCAACCACGGCTCCCACCACGCGTCGAAGAAGAGCAAGCTCCAGGTCA
AGGCGCAGCTGTCCAACCCGGGCCTGCGCGACGACGCCGAGCTGGCCGCCCGCCCGGCCTTCGACCTGGAGGGCGCCG
AGCCCACCCCCGAGCAGCGCGCCCGGGTCTTCGCCCGCAAGACGTACCGCTTCTACGAGGGCGGCGCCGTCACCCAGG
CCGACCTGCTGGGCCTGCTGGGCGCCACGGTCACCGCCGGCTACTCGCGCAAGGCGGCCGACCTGGCCCCCGCCGAAC
TCGGCCAGATCCTGCGCTGGTTCGGCCAGTACATCAGCGAGGAGCGGCTCCTGCCGAAGTACGGCTACGCCTCCCCGG
GCGCGCTGTACGCGACGCAGATGTACTTCGAGCTGGAGGGCGTCGGCGGTCTGAAGCCGGGCTACTACTACTACCAGC
CGGTCCGCCACCAGCTCGTCCTCATCAGCGAGCGCGAGGCCACCGGCAAGGCCACGGCGCAGATCCACTTCATCGGCA
AGAAGAGCGGCATCGAGCCGGTCTACAAGAACAACATCCTCGAGGTCCTGGAGATCGAGACCGGCCACATGGTCGGCC
TCTTCGAGCAGATCCTGCCGGCCTACGGCCTCGACATCCACGACCGCGCCTACGAGCCGGCCGTCAAGGACCTGCTCG
ACGTCGCCGACGAGGACTACTACCTGGGCACCTTCGAGCTGGTCCCGCACGCGGGCGCGCGCGACGACCAGGCCGAGG
TCTACGTCCAGACGCACGGCGGAAAGGTCGCCGGCCTGCCCGAGGGCCAGTACCGCTACGAGAACGGCGAGCTGACCC
GCTTCTCGGACGACATCGTCCTCAAGAAGCACGTCATCGCGATCAACCAGTCGGTGTACCAGGCCGCCAGCTTCGGCA
TCAGCGTCTACAGCCGCGCCGAGGAGGAGTGGCTGAAGTACATCACCCTCGGCAAGAAGCTCCAGCACCTGATGATGA
ACGGGCTGAACCTGGGCTTCATGTCCTCGGGCTACAGCTCCAAGACGGGCAACCCGCTGCCGGCCTCGCGCCGCATGG
ACGCCGTCCTCGGCGCCAACGGCGTCGACAGCGCCCCGATGTACTTCTTCGTCGGCGGCCGCATCAGCGACGAGCAGA
TCGGCCACGAGGGCATGCGCGAGGACAGCGTCCACATGCGCGGTCCGGCCGAGCTCATCCGCGACGACCTCGTCAGCT
TCCTCCCGGACTACATGATCCCCAACCGGGTCGTGGTCTTCGACCGGCTGCCGCTGTCCGCCAACGGCAAGATCGACG
TCAAGGCGCTCGCCGCCTCCGACCAGGTCAACGCCGAGCTCGTCGAGCGCCCCTTCGTCGCCCCGCGCACGGAGACGG
AGAAGGAGATCGCGGCGGTCTGGGAGAAGGCCCTGCGCCGCGAGAACGCCTCCGTCCAGGACGACTTCTTCGAGTCGG
GCGGCAACTCGCTGATCGCCGTCGGCCTCGTCCGCGAGCTCAACGCGCGCCTGGGCGTCTCCCTGCCGCTGCAGAGCG
TCCTGGAGTCCCCGACCATCGAGAAGCTGGCCCGCCGCCTGGAGCGCGAGGTCGCCCAGGAGTCCTCGCGCTTCGTCC
GCCTGCACGCGGAGACCGGCAAGGCCCGGCCCGTGATCTGCTGGCCGGGTCTGGGCGGCTACCCGATGAACCTGCGCA
GCCTGGCCGGCGAGATCGGCCTCGGCCGCTCGTTCTACGGCGTCCAGTCCTACGGCATCAACGAGGGCGAGACCCCGT
ACGAGACCATCACCGAGATGGCCAAGAAGGACATCGAGGCCCTCAAGGAGATCCAGCCGGCCGGCCCCTACACCCTGT
GGGGCTACTCCTTCGGCGCCCGCGTGGCCTTCGAGACCGCCTACCAGCTGGAGCAGGCGGGCGAGAAGGTGGACAACC
TCTTCCTGATCGCCCCGGGCTCCCCGAAGGTGCGCGCCGAGAACGGCAAGGTGTGGGGCCGCGAGGCGTCCTTCGCCA
ACCGCGGCTACACCACGATCCTGTTCTCGGTCTTCACCGGCACCATTTCCGGTCCGGACCTGGACCGGTGCCTGGAGA
CCGTGACGGACGAGGCCTCCTTCGCCGAGTTCATCAGCGAGCTCAAGGGAATCGACGTCGACCTCGCCCGCCGGATCA
TCTCGGTCGTGGGCCAGACGTACGAATTCGAGTACTCCTTCCACGAGCTGGCCGAGCGCACCCTCCAGGCGCCGATCA
GCATCTTCAAGGCCGTGGGCGACGACTACTCGTTCCTGGAGAACAGCAGCGGCTACTCGGCCGAGCCGCCGACGGTCA
TCGACCTCGACGCCGACCACTACAGCCTGCTGCGCGAGGACATCGGCGAGCTGGTGAAGCACATCCGCTACCTGCTCG
GCGAGTGActcgagcaccaccaccaccaccactgagatccggctgctaacaaagcccgaaaggaagctgagttggctg
ctgccaccgctgagcaataactagcataaccccttggggcctctaaacgggtcttgaggggttttttg
```

#### IndC

```
cccgcgaaattaatacgactcactataggggaattgtgagcggataacaattcccctctagaaataattttgtttaac
tttaagaaggagatatacatATGGCTAGCTGGAGCCACCCGCAGTTCGAAAAAGGCGTCGACTTAGAAAATAATATTA
CACAATGTGACTCAATCAATGATGTTTATCTTAAAGAAGAAGCAATAACATTGATGGATATGCTTGAGAGTCAACTTA
AGCACCAGGCAGATGGATATGTTGTTATTGATCAAGAAGAATCTCTCAGTTACGCTGATTTCTATTTGAGGGTGAAAG
AGATAGGGTATTGTCTGTCAGAAATTAGCTCAAAGAATTCGGTGGGTATTGGGCTTTTTTGTGATCCTTCTATAGATT
TAATTTGTGGTGCATGGGGTATTTTGTCAGCGGATAAAGCTTATTTGCCGTTATCGCCTGACTATCCAACTGAACGCC
TCAAATATATGATAGAAGATTCTGGTATTGATGTGATTTTTACGCAATCGCACTTAAAAGCACAGCTACAGGACATTG
CACCAAAATCAGTATTAATTATGACACCAGAAGATGTCGCTCTGACGATAAAAACACGAACAATAGAAGATATTCTGG
GCACAGTTCAAGTTCCTAAACCCACTAGTCTGGCTTATATTATTTATACCTCTGGTAGCACGGGTAAGCCAAAGGGAG
TGATGATTGAACATCACAGTATTGTAAATCAAATGAGATTTCTTGCAAAAGCGTTCAAATTAGGATGTCATTCCCGGA
TTTTACAGAAAACACCAATGAGTTTTGATGCGGCTCAATGGGAAATTCTAGCGCCTGCAATTGGTGGTCAAGTGATTA
TGGGTCCTTTAGGTTGCTATCGCGATCCGGATGCAATTATTAAAACCATTCTTCAGCATCAAGTAACGACTTTGCAAT
GTGTTCCTACTTTGCTACAAGCGTTACTGGATAATCCTAATTTTTTGGATTGCTTATCATTGACTCAAGTATTCAGTG
GGGGAGAAGCGCTGACAACCAAATTAGCCACGCAATTTTTGAATAGTTTTACTCACTGTGAATTAATCAATTTATATG
GCCCGACAGAATGTACGATTAATTCATCATTTTTCCGGGTGACAAATGAGACTTTGCCGAATTATCAAACCTCTATTT
CGATTGGTGCACCTGTAGATAATACCGAATACTACGTTCTTGATGATGATAGATTACCTGTGGCGGTTGGCGAAATTG
GCGAGCTTTATATTTCGGGTGCTCAATTAGCACGTGGTTATTTGCATAAACCAGAAATGACAAAAGATAAATTTATTT
GTAATCACCTTGTATCAGGAACTCAACATCAATGGTTATATCGAACGGGAGATCTGGTAACCAGAGGGGCTGATGGTA
ATACTTATTTTGTTGGTCGGGTTGATAGCCAGGTCAAATTACGAGGTTACCGTATTGAGCTTGATGAAATACGCCATG
CGATTGAAGAACATAGCTGGATAAAGACGGCGGCAATGTTAATTAAGAAGGATGCCAGAACGGGTTTCCAAAATCTCA
TCGCGTGTGTGGAATTAGATGAGAAAGAAGCTGCATTGATGGATCAAGGTAATAGTAGCTCACATCACAAATCAAAAG
CCGATAAACTACAGGTGAAAGCCCAACTTTCTAATTCTGGTTGTCGAAGTGAAGAGTTATGTGAAAATCGCCCTACAT
TCTTACTTCCTTATCAAGAAGGGGAGATAAAACAGAGAGAATATGCATTTGGACGCAAGACATATCGCTATTTTGAGG
GAACAGAAATAACGGTAGAGAAATTAAAAAAATTGCTGACAGCCACTCAATCGAATGAAATTAGCTCTTTGCCACTGA
GTCATCTAACCCTGAATGATTTCGGTTATGCATTGCGTTATTTTGGTCAGTTTACCAGCCATCAACGTTTATTGCCCA
AATATGCCTATGCTTCACCGGGTGCTCTCTATGCGACACAAATGTATTTTGAATTGCATAATGTTCTCGGTTTGGATG
CGGGGATTTACTATTATCATCCAGTGACACATAAGTTAATAAAAATTTCAACATTGAGTCGTCGGCAAATGCCAACGA
TAAAAGTGCATTTTATTGGCAAGCATGAAGCCATTGAGCCCGTTTATAAGAACAATATACAAGAAGTTCTGGAAATGG
AAGCGGGCCATATGATGGGTCTTTTTGATGACGTATTACCGGAAATTGGCTTGAGTATTGGTAAAAGTGAATATCAAG
ATGAATGTCCAGATTGGTATGATGGTGATATTCAGGATTATTATCTTGGTGCATTTGAAATATGTAGCTATGAACATG
GATTGCCGCCATTTGAGACTGATATTTATTTACAAACACATGCCCATAAAATACCTGAGATGCCGTGTGGTTTATATC
ACTTTTCTAACGGGGAATTTGTACGAATAAGTGATGATATTGTCCGAAAAAAGGATGTTATTGCGATTAATCAGCAAG
TTTATGATCGCTCCAGTTTTGGCGTGTCAATTATTCCACGCTGTGTCCCTGAATGGCATTATTATATAACACTGGGTC
GTCGGTTACATGCGTTACAAAGTAATCCATTGTATATTGGATTAATGTCATCTGGTTACAGTTCGAAGAGCAATAACG
ATTTACCTTCGGCGAAAAGGATGCGATCTATTCTCAATGCACTTGATAGACCTATGGCGGCATTTTATTTCTGCATAG
GTGGGGGTATTAGCCAAGCGCAATATATGTGTGAAGGCATGAAAGAAGATGTTGTTCATATGAAAGGGCCAGTTGAAA
TCATTAAAGATGATCTTCAACAACAACTCCCTCAATATATGATTCCAAATAAGGTATTAGTTTTCGATAAATTACCTT
TGACGGCCAATGGAAAAGTGGATTATCAATCTTTATCAGAATCTAAAGCCGTGGAGAATGTTTCAACACAGCGTCTAT
TGGTGCCATTACATACAGATACTGAAATAAGGCTTGGAAAAATTTGGATGGAAGTACTGAAATGGGATTCAGTATCTG
CCCTCGATGATTTTTTCGAAAGTGGGGGTAATTCTTTGATGGCCGTTGCAATGGTTAATAAGATCAATGCGGCCTTTA
ATATTCGTTTTCCGTTACAGATACTTTTTCAATCTCCTAATATAGCAGAATTGGCTAAGTGGATTGAACAGACAGACT
CTAAAACAATATCAAGATTAATTTTATTGAATCAGGCAAGCAAAGACCCCATTTACTGTTGGCCGGGTTTGGGCGGAT
ATCCTATGAGTTTGAGATTGCTTGCTAATAAAGTCGTTCCTGATCGGGCATTTTATGGAATACAGGCATATGGGATAA
ACGAGAGTGAAATACCGTTTTCTTCTATCCAGAGAATGGCAGAAGAGGATATTAAAGAGATAAAGAAAATACAGCCAG
AAGGGCCATATATATTGTGGGGATATTCATTTGGTGCCCGAGTAGCATTTGAAGTTGCATACCAGCTTGAACAAGCGG
GAGAAGAAGTTAACGCATTGAATTTATTGGCTCCGGGATCTCCTCATCTTGATATGAAGCAAGCGGAATATATGGATA
AAGGCGCTGAATTTACTAATCCGGCTTTTGTTAAAATACTTTTTTCTGTATTTTCTCGTTCAATCAACAGCCCAATGG
TTAAAACTTGCTTAGAACAAGTAAATAGTGAAACGACATTTATTAACTTTATATGTAGTCGTTTTAAAAACTTGGAAC
CATCATTAGTAAAACGTATCGTTAGGATTGTGACTTTGACTTATGATTTCAAGTACAGTATTGATGAGCTTTATCACA
GACACCTAAAGGCACCTATAACTATTTTCAAGGCGAATAGAGATAATGATTCATTTATCGAGGAATCGGATGTGATTT
CATCAATGTCGCCTAAAATAATTGAATTAATATCGGATCACTATCAACTGTTGGAAAGTGAAGGTGTTGCTGAGATTG
AGAAAATAATCTAActcgagcaccaccaccaccaccactgagatccggctgctaacaaagcccgaaaggaagctgagt
tggctgctgccaccgctgagcaataactagcataaccccttggggcctctaaacgggtcttgaggggttttttg
```

**Figure S8. DNA sequences encoding for BpsA and IndC.** Sequences are shown from T7 promoter to T7 terminator. Protein encoding sequences in capital letters.

**References:**

1. Cai, X., Nowak, S., Wesche, F., Bischoff, I., Kaiser, M., Fürst, R., and Bode, H. B. (2017) Entomopathogenic bacteria use multiple mechanisms for bioactive peptide library design. *Nat. Chem. 9*, 379-386.
